# Birds affected by a 2021 avian mortality event are strongly associated with supplemental feeding and ground foraging behaviors

**DOI:** 10.1101/2024.05.14.593614

**Authors:** Andrea J. Ayala, Tessa Baillargeon, Lawrence M. Gordon, Sabrina S. Greening, Lusajo Mwakibete, Joseph L. Sevigny, Caitlin E. Burrell, Christine Casey, Cindy P. Driscoll, Patrice Klein, Nicole L. Lewis, Lisa A. Murphy, Robert Poppenga, W. Kelly Thomas, Michael J. Yabsley, Roderick B. Gagne, Nicole M. Nemeth, Cristina M. Tato, David B. Needle, C. Brandon Ogbunugafor

## Abstract

In 2021, news outlets and state natural resources agencies reported a large number of avian deaths across several states in the eastern and midwestern USA. This event fomented a rapid and robust response from animal health experts from across the country. Given the clustered pattern of disease and death, an infectious etiology was rigorously investigated. No single causative pathogen was identified, leaving the cause and thus epidemiology of the mortality event unexplained. In this study, we attempted to hone in on potential causes or contributors to this event by constructing a dataset on affected birds’ life history, phylogeny, and ecology. After a preliminary analysis of these features, we developed a statistical pipeline to test two hypotheses regarding features of birds associated with the mortality event: (1) that a significant proportion of affected birds in the total sample are members of the Cornell Feederwatch list (*i.e.*, birds that consume supplemental feed, and their predators), and that (2) ground-feeding species would be significantly represented in the sample. While logistic regression models support the plausibility of the two hypotheses, they are statistically indistinguishable. We discuss the implications of these findings, propose future work, and highlight the importance of ecological and behavioral expertise in understanding epidemiological phenomena.

## 1 Introduction

During the spring and summer of 2021, the eastern coast of the United States experienced a large-scale die-off of passerines and near-passerines [1–5]. Thousands of primarily nestling and fledgling songbirds with neurological signs and periocular lesions were reported by the public to wildlife rehabilitators and state agencies [4]. Aptly dubbed the “Songbird Mortality Event,” it attracted wide-ranging media attention due to its size and spatial scope [3]. Focal geographic locations of the die-off initially began within the Mid-Atlantic region, specifically Maryland, Washington D.C. and Virginia, USA [2, 5, 6]. Soon, it was detected in the Northeast and Midwest, having spread north and westward [3]. Four peridomestic species were among those most frequently submitted to state agencies and wildlife rehabilitation centers: Common Grackles (Quiscalus quiscula), Blue Jays *(Cyanocitta cristata)*, American Robins *(Turdus migratorius)*, and European Starlings *(Sturnus vulgaris)* [2, 4, 5]. Representative carcasses were initially submitted to several veterinary diagnostic laboratories, and no consistent cause of death was determined [4]. To determine if there was a causative infectious agent of the die-off, unbiased metagenomic sequencing was performed on almost 200 specimens, including controls, representing 18 species [1]. Metagenomic analyses of RNA and DNA from affected birds utilized modern technology to explore whether there was a consistent microbial culprit that could be implicated in this outbreak. However, no single detected pathogen collectively accounted for the observed lesions [1], leaving the cause of the avian mortality event unexplained with the likelihood of an infectious pathogen lessened [7]. Given these circumstances, we undertook a data science approach to incorporating knowledge from bird ecology and natural history [8] to generate and test hypotheses for the cause of the mortality event. We utilized the same sample of birds that underwent pathogen metagenomics sampling in our data science approach [1]. Our specific objective was to identify ecological traits that linked the affected bird species to potentially inform hypotheses regarding their susceptibility to exposure to potential etiologies [8–10].

In our initial review of the sampled birds, we observed that many of the passerine and near-passerine species were those who regularly subsidized their diets with supplemental feed, i.e., food from bird feeders [11]. In addition, the reportedly affected raptor species tended to consume passerines and near-passerines as part of their natural diets [12]. This suggested to us that the etiological agent responsible for the mortality event may have been trophic by nature [13–15]. For instance, Cornell Project Feederwatch bird species are primarily urban or suburban passerines or near-passerines that frequently consume supplemental resources from feeders within their geographical ranges [16–18]. The Cornell Feederwatch Program is an initiative that utilizes data from citizen scientists regarding the species and frequency of birds at their feeders [16]. Upon reviewing the species represented by the carcasses, we were driven to further explore their natural history and functional ecology within the context of the mortality event [19–22]. Our observations led us to examine the causal role anthropogenic provisioning, i.e., bird feeders, may have played in the die-off as a potential bottom-up initiator of the observed mortality [22].

The use of bird feeders, frequently referred to in the literature as the intentional supplemental feeding of birds, is a popular activity [11, 23–26]. From a human dimensions perspective, people enjoy feeding wild birds and have reported negative impacts when guided to remove their feeders [27]. Such guidelines are usually associated with a perceived infectious disease threat or potential human-wildlife conflict [28–30]. However, the practice of supplemental feeding often generates complex ecological trade-offs among the avian communities that utilize provisioned foods [23, 31]. For instance, supplemental feeding can augment individual physiological robustness, immune function, feather growth, overall fitness, and survival, especially during periods of food scarcity, adverse conditions (e.g., cold weather), and increased energy demand (e.g., breeding, migration) [23, 26, 32–37].

On the other hand, supplemental feeding may also promote pathogen and parasite transmission by inflating the number of contacts between susceptible and infectious individuals [11, 26, 31, 37, 38]. Supplemental feeding has been reported not only to influence avian health, but also the abundance, distribution, and composition of local avian communities [32, 39]. Wildlife managers have also utilized supplemental feeding to support the conservation of endangered species, replacing diminished resources and providing foods free of environmental contaminants [24]. Despite a growing body of literature on the subject, the debate regarding the benefits versus the adverse effects of supplemental feeding to wild birds is a complex question and has yet to be resolved [27, 40–44].

Within our dataset, we also observed that a large proportion of the species represented are those that forage primarily on the ground to meet their nutritional requirements [45–47]. This is also consequential, as many ground-foraging species may be affected by contaminants or organic pesticides [48, 49]. Mechanistically, this may occur either through the probing of treated soil or the consumption of affected prey items [50–52]. Given the representation of ground-foraging species in the dataset, we could not rule out this potential hypothesis.

In this study, we address the knowledge gap regarding the role of 1) Cornell Feederwatch bird species and their avian predators, and 2) ground-foraging species in the avian mortality event of 2021. We collected metadata with respect to each species’ ecology by utilizing the Cornell All About Birds website [53]. Standardized ecological traits such as habitat, natural foods, and geographic range are published in this comprehensive online encyclopedia. Given the distribution of the data and our observations, we provide two alternative hypotheses to examine in our analyses:

1. We hypothesized Cornell Project Feederwatch species would be significantly represented within the total cases submitted to respective laboratories for pathogen sequencing.
2. We hypothesized ground-foraging species would be significantly represented in our total case sample that underwent pathogen sequencing.

To test these hypotheses, we developed a series of logistic regression models. One series of models had Feederwatch status as the dependent variable, while the second series had ground foraging species as the dependent variable. We utilized the same set of predictors to compare the most parsimonious models. We found, of the two competing hypotheses, the logistic regression model with Feederwatch status as the dependent variable had the strongest support (hypothesis one). However, the single best predictor for Feederwatch status was foraging niche – which included ground-foraging birds as one of the nominal categories (hypothesis two). The implications of these results suggest that hypothesis one (Feederwatch status) and hypothesis two (ground-foraging birds) were statistically indistinguishable from one another. Given the qualitative components of the avian mortality event – specifically the sudden onset, relatively limited duration, and the large geographical area affected, we postulate that the source of the mortality event may have been associated with ground-foraging, Feederwatch species.

## 2 Materials and Methods

### Case Acquisition

All analyses utilized data from bird carcasses that were submitted to the New Hampshire Veterinary Diagnostic Laboratory, the Wildlife Futures Program at the University of Pennsylvania, the Southeastern Cooperative Wildlife Disease Study at the University of Georgia, or the Ohio State University [1, 4]. Prior to these analyses, a large subset of these birds underwent either RNA or DNA sequencing for pathogen metagenomics [1]. Each bird categorized as a clinical case in this study demonstrated clinical signs and/or gross lesions consistent with the mortality event prior to death or euthanasia [1]. Carcasses were identified to species by wildlife rehabilitators, wildlife veterinarians, and state wildlife agency biologists who submitted the birds, and confirmed at laboratories where necropsied/sampled. Individuals that could not be speciated were not utilized in this analysis.

### Clinical Cases and Ecological Data

We performed a series of statistical analyses that examined associations between the ecological traits of the affected birds that may have influenced their susceptibility to the mortality event.

Once species identification was concluded, data for each ecological variable were extracted using the Cornell All About Birds website [53]. Thus, in using the same database for all species, we were able to maintain consistency for each variable (Tables 1 and 2). We began with phylogenetic data such as Order and Family (Table 1). Crucial to our analysis was the identification of the state from which the bird had been collected (Fig. 1). These data were provided by the individuals who submitted the carcasses for diagnostic evaluation.

**Fig. 1:**
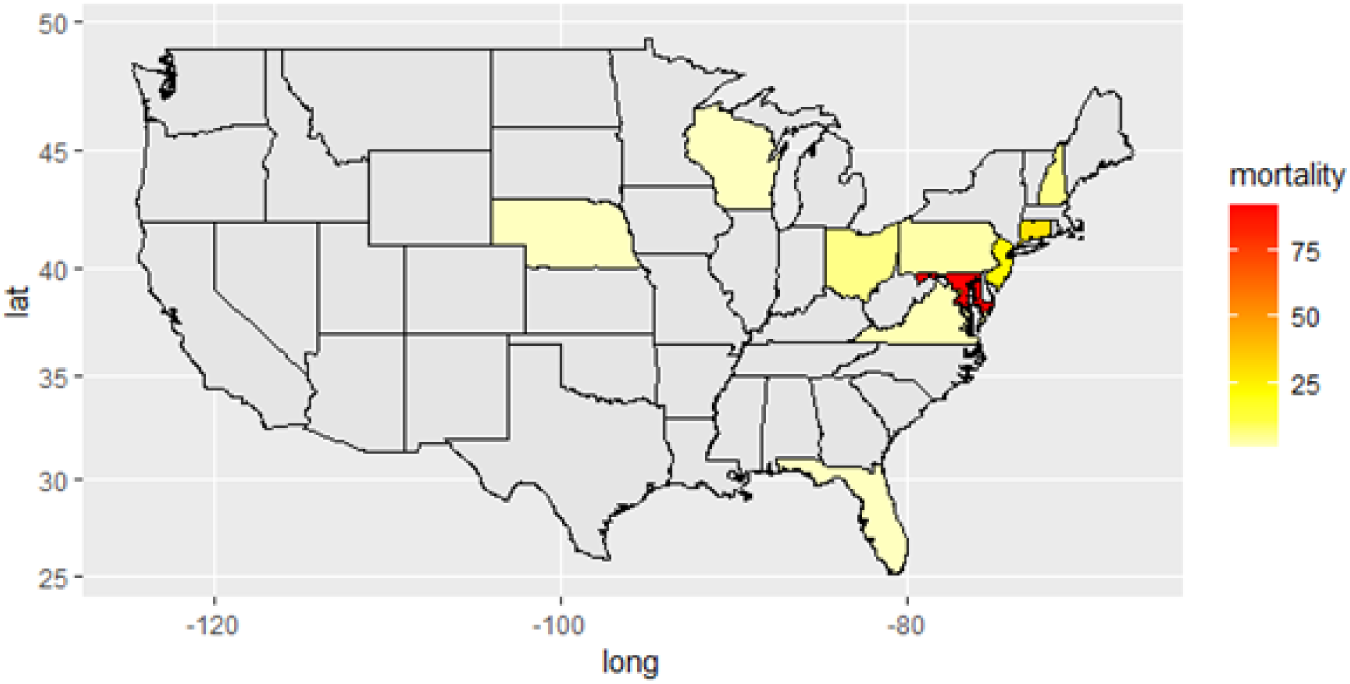
TA map of the continental United States with respect to the avian mortality event, specifically the cases that were submitted for pathogen metagenomics. States in gray had no cases reported by our dataset. Mortality (on the right side of the map) refers to the number of cases by state. Most cases were sequestered in the Mid-Atlantic region, where the mortality event was first reported (orange). Darker colors represent higher numbers of cases. Connecticut was represented by 29 cases, Washington D.C. was represented also by 29 cases, Maryland was represented by 92 cases, New Hampshire was represented by 6 cases, New Jersey was represented by 23 cases, Ohio was represented by 6 cases, Pennsylvania was represented by 3 cases, Virginia was represented by 2 cases, and fewer cases were reported in Florida, Wisconsin, and Nebraska (*n* = 1) each. It is important to note that West Virginia, Kentucky, and Arkansas also submitted carcasses, but were not a part of the sample that were submitted for pathogen metagenomics.

**Table 1:** The phylogenetic categories of affected birds are listed in this table. We identified Order and Family to determine if there were any significant relationships between the mortality event and species relatedness.

| Phylogenetic Variable | Categories | Citation |
| --- | --- | --- |
| Order | Passeriformes | [54–66] |
|  | Accipitriformes | [67, 68] |
|  | Strigiformes | [69] |
|  | Columbiformes | [70, 71] |
| Family | Corvidae | [54, 56] |
|  | Turdidae | [55] |
|  | Icteridae | [57, 65] |
|  | Accipitridae | [67, 68] |
|  | Tyrannidae | [58] |
|  | Strigidae | [69] |
|  | Sturnidae | [59] |
|  | Fringillidae | [60] |
|  | Passeridae | [61] |
|  | Columbidae | [70, 71] |
|  | Cardinalidae | [63, 64] |
|  | Mimidae | [62] |
|  | Paridae | [66] |

**Table 2:** A list of the behavioral variables and subcategories that described the affected birds.

| Behavioral Variables | Categories | Citation |
| --- | --- | --- |
| Migratory Status | Year-round | [54–56, 59–62, 66, 68–71] |
|  | Breeding | [58, 64] |
|  | Non-Breeding | [65] |
|  | Unknown (Year-round or Breeding) | NA |
| Habitat | Open Woodlands | [54, 55, 57, 58, 63, 70] |
|  | Forests | [56, 64–67, 69] |
|  | Towns | [59–62] |
| Foraging Preference | Omnivory | [54, 56, 57, 61, 62] |
|  | Insectivory | [55, 58, 59, 64–66] |
|  | Birds (Carnivory) | [67, 68] |
|  | Small Animals (Carnivory - including birds, small mammals and lizards) | [69] |
|  | Granivory | [60, 63, 70, 71] |
| Foraging Niche | Ground foraging | [54–57, 59–63, 65, 70, 71] |
|  | Aerial foraging | [67–69] |
|  | Flycatching | [58] |
|  | Foliage gleaners | [64, 66] |
| Natural Foods: Berries/Plants | Binary (Yes/No) | [54–56, 59, 62–66] |
| Natural Foods: Insects | Binary (Yes/No) | [53–58, 61–65, 67, 68, 70] |
| Natural Foods: Birds | Binary (Yes/No) | [54, 56, 57, 65, 67–69] |
| Natural Foods: Seeds | Binary (Yes/No) | [54, 56–61, 63–66, 70, 71] |
| Feederwatch Status | Binary (Yes/No) | [72] |

Using that information, we collected variables from the Cornell All About Birds website that were behaviorally relevant to each species. This included data such as whether the bird was a migrant or year-round resident of the state from which it was collected, as well as its primary habitat, i.e., open woodlands, forested areas, or towns. Primary feeding behaviors were characterized by the Cornell All About Birds site as either omnivory, insectivory, granivory, or predators of birds or predators of animals [53]. Feeding niche was categorized by Cornell as ground-foraging, aerial feeding, flycatching, or foliage gleaning (Table 2) [53]. We also noted natural foods that were part of the species’ diet due to potential overlaps in feeding preferences, such as berries/plants, insects, other birds, and seeds (Table 2) [53].

### Statistical Analyses of Ecological Variables

All statistical analyses were performed using R version 4.2.2 [73]. Prior to all analyses, individual birds identified as ‘controls’ were removed from the dataset [1, 4]. A schematic detailing our statistical workflow is available below (Figure 2).

**Fig. 2:**
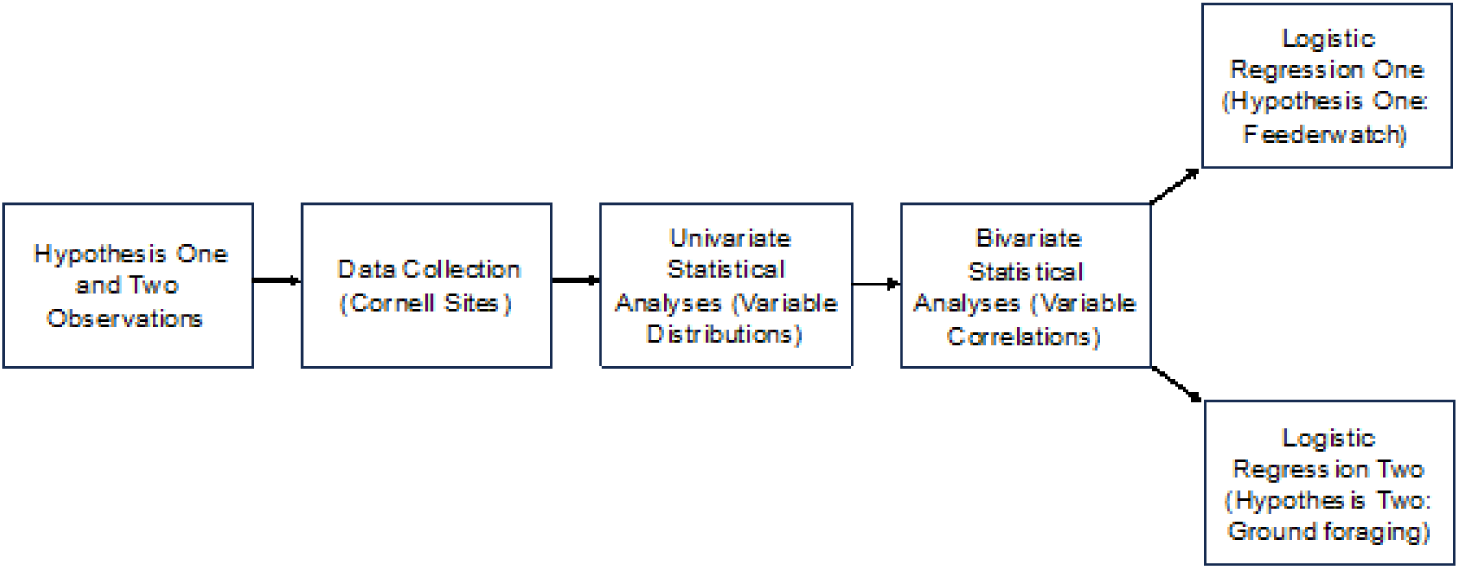
A detailed schematic of our statistical workflow. We began our analyses with observations of our hypotheses (step 1), and then collected the data at the Cornell websites (step 2). We then performed univariate analyses of our variables (step 3), followed by bivariate analyses to determine whether our variables were autocorrelated (step 4). Finally, we tested our hypotheses using two series of Logistic Regression models, one each that tested a single hypothesis (step 5).

We assessed the data distributions for the number of cases associated with the phylogenetic variables Order and Family, and the ecological variables Migratory Status, Foraging Preference, and Foraging Niche using the Cullen and Frey distribution plot [74, 75]. Habitat was not assessed using the Cullen and Frey distribution plot due to the limited number of categories. The purpose of this test was to assess the amount of skewness and kurtosis associated with the cases for these variables, and determine if they fell within a Gaussian, i.e., normal distribution [76]. Binary variables are represented by the logistic distribution [77], and thus were not assessed using the Cullen and Frey distribution plot.

We also performed Chi-Square Goodness of Fit tests for the same variables [78]; however, we were able to include the binary variables Feederwatch Status, Natural Foods: Birds, Natural Foods: Insects, Natural Foods: Non-feeder Seeds, and Natural Foods: Berries and Plants. Family was not assessed as it was represented by 13 categories, and we had insufficient sample size power to make meaningful inferences. The purpose of the Chi-Square Goodness of Fit test was to determine whether the number of cases represented by these variables were uniformly distributed [79]. When necessary, follow-up multiple comparison Chi-Square tests were performed to determine which sub-categories demonstrated statistical significance. Bonferroni corrections to the significance value of *p* = 0.05 were applied to these post-hoc tests [80].

Given the number of ecological variables we examined, we also had to address the possibility of multicollinearity across the independent variables within our logistic models [81]. We determined the degree of correlation between our ecological predictors, i.e., independent variables, which were all categorical. This was achieved by performing a series of Chi-Square Tests of Independence and setting our significance value to *p* = 0.05 [82]. We also provided the Cramer’s V result, which is a statistical measure of the association between two categorical variables that are not represented by standard 2 x 2 contingency tables. The values for Cramer’s V range from zero, which indicates no association, and one, which indicates a perfect association [83].

As our first dependent variable was Feederwatch Status, we assessed whether the independent variables in this study were predictive of Feederwatch Status. We utilized hypothesis-driven logistic regression, followed by likelihood ratio tests, to identify which phylogenetic, behavioral and/or natural food traits would be predictive of Feederwatch Status. This approach was used to assess whether susceptibility to involvement in this mortality event was associated with accessing supplemental feed.

Given that our second dependent variable was foraging niche, specifically ground-foraging, we also assessed whether the remaining independent variables in our study were predictive of ground-feeding classification. To conduct a comparative approach, we also utilized logistic regression to identify which phylogenetic, behavioral, and/or natural food traits would be predictive of foraging niche, with respect to ground-foraging, as a binary variable (yes/no). This statistical approach could suggest that birds that consume foods primarily from the ground would have a greater probability of exposure to the etiological agent, such as would occur with a large-scale contaminant or pesticide [48, 84].

## 3 Results

### Phylogeny

A total of 197 avian carcasses underwent high-throughput metagenomic next-generation sequencing for the presence of a single RNA or DNA pathogen that would account for the mortality event [1]. This includes carcasses that were utilized as non-affected controls (*n* = 8 controls), specifically passerines that did not develop clinical signs or pathology consistent with the mortality event [1, 4]. That total also included carcasses from which species identification could not be ascertained with full confidence (*n* = 2 unknowns); these unknowns included a blackbird spp. and a sparrow spp. The removal of controls and unknowns left a total of 187 avian carcasses representing 18 species (*n* = 18 species) for analysis with respect to our variables of interest (Figure 1). These species represented a total of 13 taxonomic Families (Figure 3). The families were representative of just four Orders, the Accipitriformes (*n* = 6 Accipiters, 3.2% of cases), the Columbiformes (*n* = 3 Columbids, 1.6% of cases), the Passeriformes (*n* = 177 Passerines, 94.7% of cases), and the Strigiformes (*n* = 1 Strigid, 0.53% of cases). In terms of distribution, the Cullen and Frey plot suggested that the Beta distribution was the most appropriate with which to categorize the case-associated data for the variable Order, with an estimated skewness of 1.996649 and estimated kurtosis of 6.988749. A similar result was found for the variable cases by Family, whereby the Cullen and Frey plot also suggested the data originated in the Beta distribution, with an estimated skewness of 3.201349 and an estimated kurtosis of 13.77649.

**Fig. 3:**
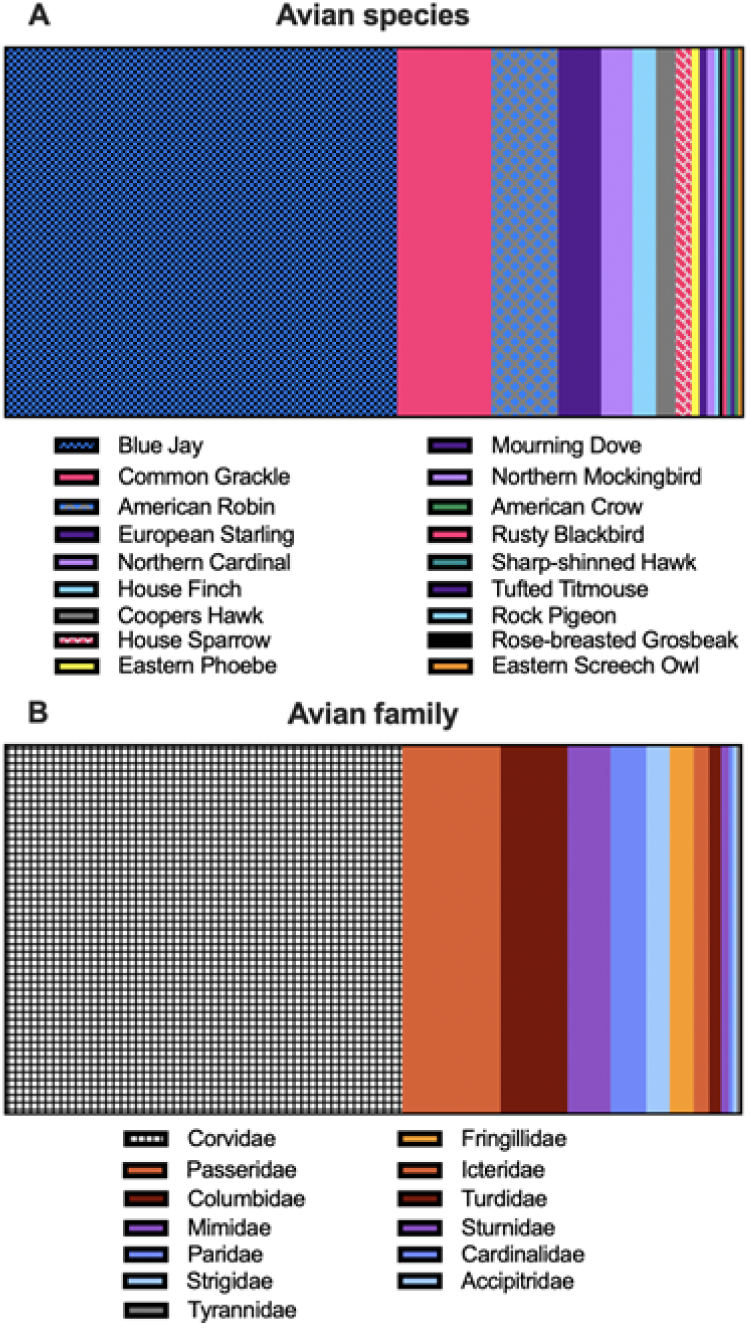
Affected birds by species. A) Eighteen confirmed species in the study, parsed out by the number of clinical cases they represent. Blue Jays (*Cyanocitta cristata*) had the highest case count, with n = 100 cases, followed by Common Grackles (*Quiscalus quiscula*), with n = 24 cases. American Robins (*Turdus migratorius*) were represented by n = 17 cases, and European Starlings had n = 11 cases. Northern Cardinals (*Cardinalis cardinalis*) were represented by n =8 cases, followed by House Finches (*Haemorhous mexicanus*) with n = 6 cases. Five Cooper’s Hawks (*Accipiter cooperii*), n = 5 cases, were represented in the sample, and four House Sparrows (*Passer domesticus*), n = 4 cases were represented. Two each of Mourning Doves (*Zenaida macroura*) n = 2, and Northern Mockingbirds (*Mimus polyglottos*), n = 2 were submitted as cases. One each of an American Crow (*Corvus brachyrhynchos*), Rusty Blackbird (*Euphagus carolinus*), Sharp-shinned Hawk (*Accipiter striatus*), Eastern Phoebe (*Sayornis phoebe*), Rock Pigeon (*Columba livia)*, Tufted Titmouse (*Baeolophus bicolor*), Eastern Screech Owl (*Megascops asio*), and Rose-breasted Grosbeak (*Pheucticus ludovicianus*), n = 1 case, had cases in our sample set. B) The number of cases as distributed by avian family. The Corvidae (Blue Jays, and an American Crow) had the largest number of cases, while the Paridae (*Tufted Titmouse*), Strigidae (Eastern Screech Owl), and Tyrannidae (*Eastern Phoebe*) had the fewest cases with one each.

The Chi-Square Goodness of Fit Test for uniformity of the cases across Orders was statistically significant (*χ*^2^ = 484.12, df = 3, p-value *<* 0.0001), thus indicating a non-uniform distribution. Given that there were four Orders, we performed a Bonferroni correction to identify where the significance lay (*p* = 0.0083). We found that cases among passerines (Order Passeriformes) were significantly over-represented in comparison to the other Orders (Table A1).

### Project Feederwatch Status

As our first dependent variable, we assessed the number of cases associated with the binary variable Feederwatch status. A ‘yes’ indicated that those cases were associated with a Project Feederwatch species. A ‘no’ indicated that the respective cases were not associated with a Project Feederwatch species. A total of 177 cases were associated with Feederwatch species (*n* = 177 cases, 94.7%), while ten cases were not associated with Feederwatch species (*n* = 10 cases, 5.3%). The Chi-Square Goodness of Fit Test for uniformity of cases across Feederwatch status was statistically significant, suggestive of a non-uniform distribution (*χ*^2^ = 149.14, df = 1, p-value *<* 0.0001).

### Foraging Niche

We utilized foraging niche as our second dependent variable. We first examined four categories of foraging niches, or how birds prefer to feed, representative of the cases associated with the species in our sample. The highly maneuverable aerial foragers were associated with 7 cases (*n* = 7 cases, 3.7%). Flycatching birds that ‘sally’ were associated with a single case (*n* = 1, 0.53%), while foliage gleaners were associated with two cases (*n* = 2, 1.1%). Ground foragers were associated with 177 total cases (*n* = 177 cases, 96.7%). Our Chi-Square Goodness of Fit test was statistically significant (*χ*^2^ = 484.29, df = 3, p-value *<* 0.0001), suggesting the data were not uniformly distributed. Upon following up with a series of Chi-Square tests, we applied a Bonferroni correction (p-value = 0.0083) (Table A2). In order to utilize this variable as the response variable in a logistic regression, we subsetted the category ground-foraging from foraging niche. We subsequently converted the variable ground-foraging into a stand-alone binary variable (yes/no). The results of the Cullen and Frey plot suggested that the distribution for the cases associated with the variable Foraging Niche fell into the Beta distribution, with an estimated skewness of 1.99456 and an estimated kurtosis of 6.981411.

### Migratory Status

Migratory status served as an independent variable. We assessed the number of cases associated with each migratory status with respect to the state in which each bird was collected. This was parsed out according to the categories inclusive of year-round residents, breeding birds only, and non-breeding birds, or those for which range maps did not discern at the state level (unknowns).

We found that the majority of cases were associated with year-round resident birds (*n* = 183 cases, 97.9%), while breeding birds (*n* = 2 cases, 1.1%), non-breeding birds (*n* = 1 case, 0.53%) and unknowns (*n* = 1 case, 0.53%) were fewer in number.

The Chi-Square Goodness of Fit Test for uniformity of the cases across migratory status was statistically significant, suggestive of a non-uniform distribution (*χ*^2^ = 529.47, df = 3, p-value *<* 0.0001). Given that there were four groups, we performed a Bonferroni correction to confirm significance (*p* = 0.0083). We found that cases represented by year-round residents were significantly over-represented in comparison to the other groups (Table A3). The Cullen and Frey distribution plot suggested that the cases associated with these categories originated in the Beta distribution, with an estimated skewness of 1.999839 and kurtosis of 6.999459.

### Foraging Preference

We assessed five categories of foraging preferences, or primary foods, for the cases associated with the species in our sample. Birds that consume primarily other birds were associated with six cases (*n* = 6 cases, 3.2%). Birds that consume primarily insects, also known as insectivores, were represented by 32 cases (*n* = 32 cases, 17.1%). Omnivorous birds were represented by 131 cases (*n* = 131 cases, 70.1%). Granivores, or birds that primarily consume seeds, were represented by 17 cases (*n* = 17 cases, 9.0%), and birds that primarily consume small vertebrates were represented by one case (*n* = 1 case, 0.53%).

The Chi-Square Goodness of Fit Test for uniformity of the cases across foraging preferences was statistically significant (*χ*^2^ = 307.95, df = 4, p-value *<* 0.0001). Given that there were five foraging preference categories, we performed a Bonferroni correction to identify where the significance lay (p-value = 0.005). We found that cases represented by omnivorous birds were significantly over-represented in comparison to the other foraging preferences (Table A4). The Cullen and Frey distribution plot suggested that the cases represented by each of these foraging preferences were represented by the Beta distribution, with an estimated skewness of 1.974617 and an estimated kurtosis of 7.010337.

### Habitat

We assessed three categories of species habitat preferences with respect to the cases in our sample. Birds that inhabit forests represented 110 cases (*n* = 110 cases, 58.8%), while open woodland dwellers represented 53 cases (*n* = 53, 28.3%) and town birds represented 24 cases (*n* = 24, 12.8%). The Chi-Square Goodness of Fit test revealed that the distribution of cases according to habitat preference were not uniform in nature (*χ*^2^ = 61.422, df = 2, p-value *<* 0.0001). The applied Bonferroni correction for follow-up Chi-Square testing provided a p-value*of* 0.0166 (Table A5). No Cullen and Frey distribution plot could be run for this variable as it did not have a minimum of four categories associated with cases.

### Natural Foods: Birds

We next examined the number of cases associated with birds in our representative sample for whom the consumption of other birds is part of their diet, albeit not specifically their primary source of nutrition. As a binary (yes/no) characteristic, we assessed this variable using the Chi-Square Goodness of Fit test to determine the uniformity of cases across species. Fifty-five (*n* = 55 cases, 29.4%) were not associated with birds that either primarily or facultatively consume other birds, while 132 (*n* = 132 cases, 70.6%) were associated with birds that facultatively or primarily consume other birds. This result was statistically significant (*χ*^2^ = 31.706, df = 1, p-value *<* 0.0001).

### Natural Foods: Insects

We assessed the number of cases associated with birds in our sample that consume insects, however, with the caveat that this is not their primary source of nutrition for the adults of these species. As a binary (yes/no) characteristic, we assessed this variable using the Chi-Square Goodness of Fit test to determine the uniformity of cases. Thirteen (*n* = 13 cases, 6.95%) were not associated with the consumption of insects, while 174 (*n* = 174 cases, 93.04%) were associated with the consumption of insects. This result was statistically significant (*χ*^2^ = 138.61, df = 1, p-value *<* 0.0001).

### Natural Foods: Seeds

We delved into the number of cases associated with birds in our representative sample for whom the consumption of naturally occurring seeds is a part of their diet, although not their primary diet. As a binary (yes/no) characteristic, we assessed this variable using the Chi-Square Goodness of Fit test to determine the uniformity of cases across species. Twenty-six (*n* = 26 cases, 13.9%) were not associated with the consumption of naturally occurring seeds, while 161 (*n* = 161 cases, 86.1%) did consume seeds. This result was statistically significant (*χ*^2^ = 97.46, df = 1, p-value *<* 0.0001).

### Natural Foods: Berries and Plants

We next examined the number of cases associated with the consumption of plant matter such as berries and plants in our representative sample. This is not specific to birds for whom herbivory is their primary source of nutrition. As a binary (yes/no) characteristic, we assessed this variable using the Chi-Square Goodness of Fit test to determine the uniformity of cases across species. One hundred and twenty-one birds (*n* = 121 cases, 64.7%) were not associated with the consumption of berries or plant matter, while 66 (*n* = 66 cases, 35.29%) do consume berries or plant matter. This result was statistically significant (*χ*^2^ = 16.176, df = 1, p-value *<* 0.0001).

### Bivariate Chi-Square Tests of Association

Across multiple taxa that are contingent upon natural, ecological conditions, life history traits often covary with one another [85–88]. Therefore, in order to test our two hypotheses (Feederwatch status versus ground-foraging birds), we first needed to identify which variables may have been subjected to multicollinearity in our larger logistic models [82, 89]. In light of that, we performed a series of Chi-Square Tests of Independence between our variables to identify those that were highly correlated to one another (Table A6).

### Logistic Regression Model for Hypothesis One: Feederwatch species status

Given that Project Feederwatch species accounted for 177 cases or 96.7% of the evaluated cases, we utilized a series of logistic regression models to identify the phylogenetic or behavioral variables predictive of Project Feederwatch species status. We sought to identify the most parsimonious model of natural history traits that accounted for a case having the potential to visit supplemental feeders during the mortality event. In Table 3, we list the logistic models we examined to account for the variance in Project Feederwatch species status as our dependent variable.

**Table 3:**
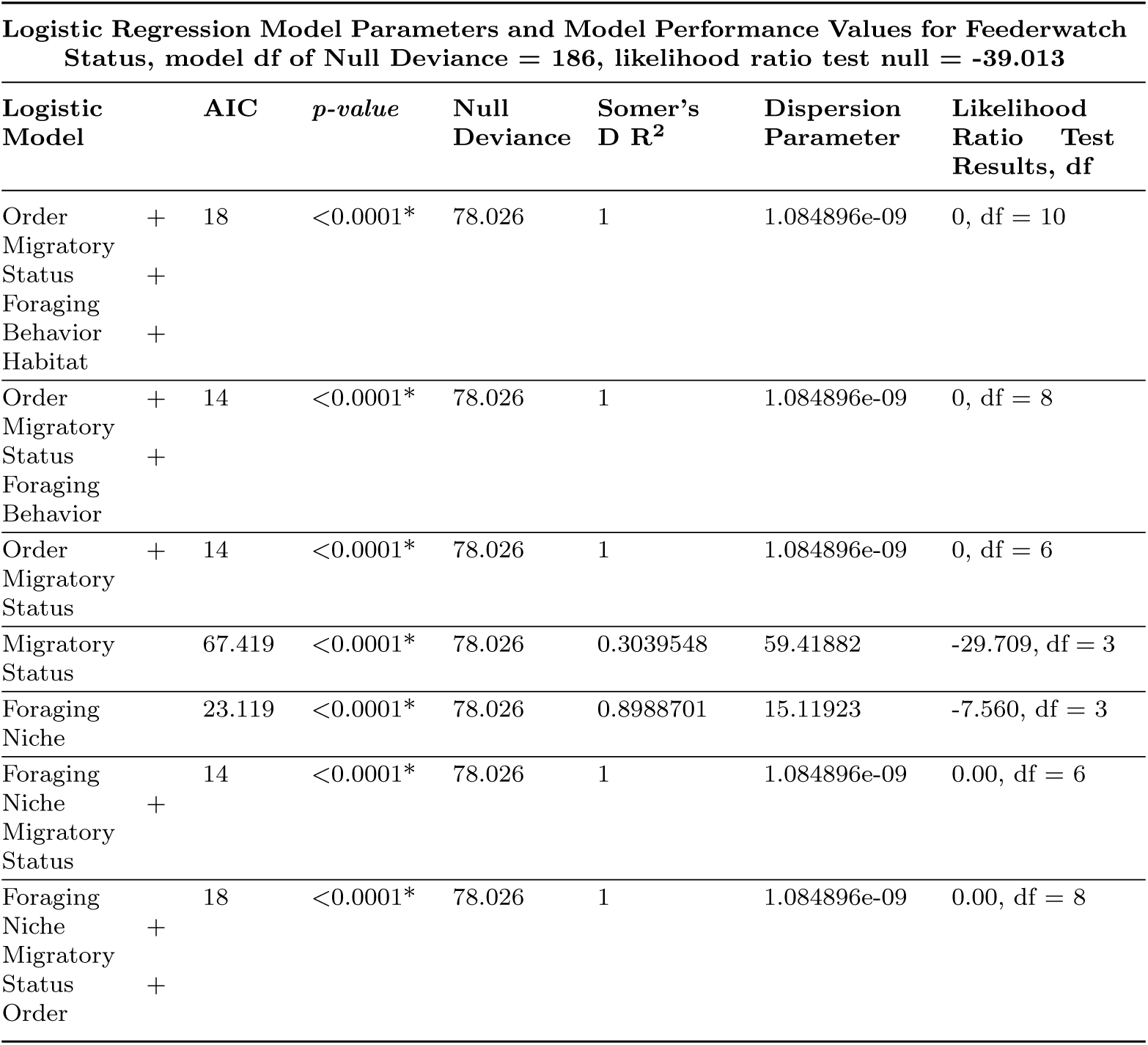
A summary of the logistic regression models that were analyzed, with Feederwatch status as the dependent variable. The AIC, or Akaike’s Information Criterion scores models based on model fit, in which a lower score is more predictive of a better model fit. The null deviance represents the measure of variance the response variable, while the dispersion parameter is a measure of overdispersion of the model - in which model parameters are over inflated with respect to their contribution to the model. Finally, the likelihood ratio tests compare the goodness of fit of multiple nested models. The p-values for each model were computed against the null model, which is represented by the default intercept of 1. An asterisk (*) denotes that the p-value is statistically significant.

The data suggested the best model accounted for up to 89.9% of the total variance, as all of the models that reported a value of 1 were overfitted [90, 91]. The most parsimonious model was generated by the independent variable Foraging Niche, with an AIC value of 23.119 (Table 3) [92], and a statistically significant model p-value of *<* 0.0001, with three total degrees of freedom (df = 3). In addition, we utilized the Somer’s D test statistic to indicate model fit. In logistic regression, this test provides an estimate of the correlation of the observed binary response variable and the predicted probabilities [93]. Our model fit was 0.8988, which indicates that up to 89.9% of the model variance was accounted for.

### Logistic Regression Model for Hypothesis Two: Ground foraging

Within the Foraging Niche covariate, the subset of ground-foraging species accounted for 177 total cases (*n* = 177 cases, 96.7%) of the clinical cases. Thus, we again utilized a series of logistic regression models to identify the phylogenetic or behavioral variables predictive of ground-foraging species status as a stand-alone binary variable (yes/no). We utilized the logistic regression approach in this hypothesis so the results could be comparable to the logistic modeling utilized in the first hypothesis (Project Feederwatch species status). Thus, we sought to identify the most parsimonious model of natural history traits that may have accounted for a case being associated with ground foraging characteristics just prior to and during the mortality event. In Table 4, we list the logistic models we examined to account for the variance with ground-foraging species status as our dependent variable.

**Table 4:** A summary of the logistic regression models that were analyzed, with ground-foraging status as the dependent variable. The p-values for each model were again computed against the null model, which is represented by the default intercept of 1. An asterisk (*) denotes that the p-value is statistically significant.

| Logistic Regression Model Parameters and Model Performance Values, model df of Null Deviance for Foraging Niche, with respect to Ground foraging = 186, likelihood ratio test null = -39.013 |  |  |  |  |  |  |  |
| --- | --- | --- | --- | --- | --- | --- | --- |
| Logistic Model | AIC | p-value | Null Deviance | Somer's D R<sup>2</sup> | Dispersion Parameter | Likelihood Ratio Test Results, df |  |
| Order +<br>Migratory Status +<br>Foraging Behavior +<br>Habitat | 22 | $<0.0001^*$ | 78.026 | 1 | 1.855982e-09 | 0, df = 10 | |
| Order +<br>Migratory Status +<br>Foraging Behavior | 28.7 | $<0.0001^*$ | 78.026 | 0.984 | 8.699706 | -4.350, df = 9 | |
| Order +<br>Migratory Status | 26.301 | $<0.0001^*$ | 78.026 | 0.902 | 12.30079 | -6.150, df = 6 | |
| Migratory Status | 73.16 | 0.006423* | 78.026 | 0.20 | 65.72578 | -32.863, df = 3 |  |

The data suggested that the best model accounted for up to 90.2% of the total variance, as all of the models that reported a value of 1 were overfitted [90, 91]. The most parsimonious model was generated by the independent variables Order and Migratory Status, with an AIC value of 26.301 (Table 4), and a statistically significant model p-value of *<* 0.0001, with six total degrees of freedom (df = 6). In addition, we again utilized the Somer’s D test statistic to indicate model fit [86]. Our model fit was 0.902, which indicates that up to 90.2% of the model variance is accounted for.

## 4 Discussion

In this study, we documented a statistically significant association between the evaluated birds [1, 4] that were part of a widespread mortality event with Feederwatch status [72]. We also assessed the associations between cases with Foraging Niche [94], leading to a similarly statistically significant association.

Furthermore, in our logistic regression model for hypothesis one (Feederwatch status), we found that the significant predictor of Feederwatch status was the variable Foraging Niche [50]. The logistic regression for hypothesis two, which had a higher AIC value than the logistic regression for hypothesis one, suggested that the most significant predictors for Foraging Niche were Order and Migratory Status. Since the same data were used for both hypotheses, the AIC values for each logistic regression model were comparable.

Based on similar *R*^2^ values and a lower AIC [92, 95], the data suggest that the logistic regression for model one had the strongest support. However, the strongest predictor for Feederwatch Status was Foraging Niche. This suggests that hypothesis one (Feederwatch species status) and hypothesis two (ground foraging) are statistically indistinguishable from one another, as ground-foraging birds made up 96.7% of the cases represented in Foraging Niche. Based on the temporal and spatial movement of the mortality event, and the toxicological tests that were performed, however, it seems less likely that a mycotoxin, contaminant or pesticide was the causative agent. We suggest this because the etiological agent would have required the same contaminant or pesticide to be applied to the Northeast and Midwest on a large-scale basis in a short amount of time (summer 2021), causing the observed lesions and mortality in songbirds and their avian predators. Given that no other ground foraging birds were affected, i.e., shorebirds, or herons, hypothesis two lacks qualitative support.

However, it should be noted that a potential etiology for the periocular swelling and irritation may have been due to a spray pesticide directed against the spring and early summer arising insects, which included the Brood X cicadas. Overall, the logistic model for our second hypothesis supports that sedentary (year-round) residents of the Order Passeriformes were statistically represented in our sample of clinical cases. This may also be consistent with ground foraging Feederwatch Status birds.

An intriguing feature of both hypotheses was the relatively large representation of Blue Jays (*n* = 100 cases), American Robins (*n* = 17 cases), European Starlings (*n* = 11) and Northern Cardinals (*n* = 9). These species have widely variable natural histories; however, they are also Feederwatch status birds who tend towards feeding on the ground [88–90]. The species that did not fall into the categories of Feederwatch birds were predominantly the avian predators, i.e., Eastern Screech Owl, Sharp-shinned Hawk, and the Cooper’s Hawks. The two passerines (Order Passeriformes) that did not fall into the Feederwatch category were the Eastern Phoebe, which is often seen sallying for insects near or at feeders, and the Rose-breasted Grosbeak, which does occasionally visit feeders, especially during migration [35, 96]. With respect to the ground-foraging category, it should also be noted that of the two passerines that did not fall into the Cornell ground-foraging designation, Eastern Phoebes are noted for their perch-to-ground sallies [97]. The second species, Tufted Titmice, will often forage in the leaf litter [98]. With respect to European Starlings, invasive species are not always admitted to rehabilitation centers, so they may have been under-represented in the sample.

Lastly, most of the submitted avian carcasses consisted of nestling and fledgling age classes of birds, which has implications for both hypotheses. The rationale is that in hypothesis one (Feederwatch status), many songbirds often supplement their nestlings with food from feeders if available. However, assessment of impacts of supplemental feeding requires further observation and/or experimentation with Feederwatch birds to determine the extent to which supplemental feed of various types may be regurgitated to nestlings. In addition, fledglings are also often drawn to feeders themselves or by their parents, as it may be easier to consume supplemental feed than to ‘hunt’ for food. With regard to hypothesis two, ground foragers may have ingested insects that were coated with or had themselves consumed contaminants, and then fed those to their young. Coupled with potential immunosuppression from even small quantities of mycotoxin or other contaminants that were potentially below the limit of detection of the toxicological tests (e.g., originating from spoiled feed or pesticide sprays), these would likely have had more adverse health impacts in younger birds [25, 99–102]. Such outbreaks are often multifactorial, and one possibility is that these events could have initiated a cascading dysbiosis that allowed common avian pathogens, such as those reported by Mwakibete et al. 2024 [1] to overwhelm the affected birds.

### Study Limitations

There are inherent limitations in wild bird diagnostic data [103, 104]. These include human observation and reporting biases, often poor-quality samples due to extended postmortem intervals and multiple freeze-thaws, as well as varied and inconsistent diagnostic approaches. In addition, case inclusion criteria were challenging to establish in this mortality event, and thus may have been inconsistent across labs and diagnostic services [4].

From the standpoint of our samples, the number of representative birds per taxa varied widely. For instance, birds that primarily consume other birds did not constitute a significant number of individuals in our dataset. However, the significance of their representation among the clinical cases is suggestive of a potential trophic mechanism behind the mortality event. In addition, birds such as Corvids and Icterids, which accounted for 125 of the 187 clinical cases, will facultatively consume nestlings or even carrion [105–107]. Surprisingly, Feederwatch status was not strongly associated with primary habitat, given that we would have expected more ‘urban’ or ‘suburban’ birds to be represented. However, birds of the same species will often partition out their habitat preferences, often inhabiting varying gradients of urbanization [108–110]. The answer to the question regarding how an individual member of a species chooses their habitat and/or territories is also one that remains elusive [110].

We must also address the representation of Blue Jays in our dataset (*n* = 100, 53%), which may have been overrepresented due to size, visibility, and carcass persistence as opposed to the smaller birds represented by fewer samples [104, 111]. Given the sheer number of Blue Jays in relation to the other species in the dataset, they likely influenced the patterns that were elicited from the logistic models. However, with few exceptions, many of the species we discussed were ground foragers and/or Feederwatch birds. For instance, birds that fell into both the categories of ‘ground-foraging’ and Feederwatch included American Robins (*n* = 17, 9.1%), Common Grackles (*n* = 24, 12.8%), European Starlings (*n* = 11, 5.8%), and Northern Cardinals (*n* = 8, 4.3%), which represent another 32% of the dataset. Thus, Blue Jays are subject to much of the variance, but this variance is also predicated upon the timing and ending of the event, and how birds were reported to and/or submitted to state and federal agencies and wildlife rehabilitators. Given the somewhat rapid dwindling of the mortality event and the biases associated with case submission, it is imperative to note that it remains unknown whether the proportions of the birds in this dataset were relational to the mortality event.

As these cases were reported to wildlife rehabilitators and/or federal and state agencies, it was also possible that the cases in this dataset were more visible to the public [19, 103]. For instance, Feederwatch species may have been more visible to members of the public, which may have instigated their reports. Thus, their collection may or may not have been associated with the species’ primary habitat as delineated by the Cornell All About Birds site [112]. In addition, the two species that were categorized in non-breeding habitat were also migratory [113], which may have affected their visibility, presenting a detection bias [114]. All of these points suggest our data consisted of a non-random convenience sample, a type of non-probabilistic sampling which is subject to both selection and sampling bias [104]. However, this method of sampling (i.e., passive surveillance) is common in wildlife mortality events [115], which often are unexpected and rely on limited resources and logistical challenges [116–118].

### Conclusions

We found strong associations between our representative sample of clinical cases and Feederwatch status, i.e., the consumption of supplemental feed in concert with ground foraging. Furthermore, that association was further correlated with the Order Passeriformes (passerines, or songbirds), and migratory status, specifically sedentary, or year-round resident birds. Our analysis supports the need for further research into the etiology and other contributing factors in the present mortality event and provides a framework for analyses of future wild bird mortality events.

Our findings do not confirm nor refute any mechanistic explanation for the cause of the mortality event. While an infectious etiology remains a possible cause or contributor to this outbreak, currently available data suggest involvement of a recognized wild bird pathogen is unlikely [1, 4]. Although the temporal and spatial aspects of the mortality event do not appear to be consistent with a toxic etiology, the role that toxicants such as mycotoxins or pesticides might play in future events needs further and consistent investigation, particularly consideration of the potential role of mycotoxins—associated with supplemental feed [119, 120]—that may have contributed to this mortality event [121]. In light of this, animal health professionals should consider (and test for) both infectious causes with non-traditional routes (e.g., fomite), and potential non-infectious causes early in the shape of large-scale die-offs that resemble infectious etiologies. Exploring potential etiologies more broadly in wildlife studies is economically costly in a field that is already challenged by limited resources [4, 122–124]. However, such considerations may help mobilize the necessary expertise and resources that would facilitate rapid and more accurate testing of affected animals for chemical, pharmacological, or biotoxin-related agents (in addition to infectious agents).

Further, our approach highlights the necessity of organismal biology and animal ecological perspectives in helping to disentangle complex epidemiological phenomena.

## Supplementary information

Tables A1-A6 contain the products of additional statistical analyses as outlined in the main text.

## Acknowledgements

The authors would like to thank extended members of the Songbird Mortality Event team, located across multiple institutions, including: The New Hampshire Veterinary Diagnostic Laboratory, the Wildlife Futures Program and PADLS New Bolton Center at the University of Pennsylvania, the Southeastern Cooperative Wildlife Disease Study at the University of Georgia, Yale University, the U.S. Geological Survey National Wildlife Health Center, and the Ohio State University. We are grateful to SCWDS staff and students, and member state wildlife management agencies and federal wildlife agency partners, including the U.S. Fish and Wildlife Service National Wildlife Refuge System and the U.S. Geological Survey Ecosystems Mission Area, for continued financial support of the SCWDS Research and Diagnostic Service. The authors would also like to thank Second Chance Wildlife Center, Tri-State Bird Rescue and Research, wildlife rehabilitators in Maryland, The Connecticut Department of Energy and Environmental Protection, and the many wildlife biologists, veterinarians and rehabilitators who dealt with this morbidity/mortality event and provided individual birds to this study. The authors would like to thank members of the Ogbunu, Almagro-Moreno, Ezenwa, and Prum labs, as well as the Yale Statistics Clinic for helpful discussions on topics related to this manuscript. Special thanks to Ketty Kabengele for helpful editorial insight, and to Valerie Shearn-Bochsler for helpful interactions.

## Declarations

### Ethics approval and consent to participate

Not applicable.

### Data availability

Data and code can be found on GitHub: https://github.com/OgPlexus/songbird1.

### Author contribution

Conceptualization: AJA and CBO. Data collection: AJA and TB. Data analysis: AJA and CBO. Data visualization: AJA and CBO. Data interpretation: AJA, LMG, SSG, LM, JLS, EPB, CEB, CC, CPD, PK, NLL, LAM, RP, WKT, MJY, RBG, NMN, CMT, DBN, CBO. Writing-original draft: AJA and CBO. Writing-revision and editing: AJA, LMG, SSG, LM, JLS, EPB, VSB, CEB, CC, CPD, PK, NLL, LAM, RP, WKT, MJY, RBG, NMN, CMT, DBN, CBO. Funding Acquisition: AJA, DBN, RBG, CMT, NMN, MJY, SSG, WKT, CBO. Supervision: AJA, RBG, NMN, CMT, DBN, CBO.

## Appendix A Supplementary information

Tables A1-A6 contain the products of additional statistical analyses as outlined in the main text.

**Table A1:** Statistical results of the follow-up Bonferroni tests with the number of cases associated with each avian Order. An asterisk (*) denotes that the p-value is statistically significant.

| Goodness of Fit Test Bonferroni Corrections: Distribution of Cases Across Orders, df = 1 |  |  |  |
| --- | --- | --- | --- |
| Variable One | Variable Two | $\chi^2$ value | p-value |
| Passeriformes | Accipitriformes | 159.79 | <0.0001* |
| Passeriformes | Columbiformes | 168.2 | <0.0001* |
| Passeriformes | Strigiformes | 174.02 | <0.0001* |
| Accipitriformes | Columbiformes | 1 | 0.3173 |
| Accipitriformes | Strigiformes | 3.5714 | 0.05878 |
| Columbiformes | Strigiformes | 1 | 0.3173 |

**Table A2:** Statistical results of the follow-up Bonferroni tests with the number of cases associated with each foraging niche. An asterisk (*) denotes that the p-value is statistically significant.

| Goodness of Fit Test Bonferroni Corrections: Distribution of Cases Across Foraging Niche, df = 1 |  |  |  |
| --- | --- | --- | --- |
| Variable One | Variable Two | $\chi^2$ value | p-value |
| Aerial | Flycatching | 4.5 | 0.03389 |
| Aerial | Foliage gleaning | 2.8 | 0.09558 |
| Aerial | Ground forager | 157.07 | <0.0001* |
| Flycatching | Foliage gleaning | 0.33333 | 0.5637 |
| Flycatching | Ground forager | 174.02 | <0.0001* |
| Foliage gleaning | Ground forager | 171.09 | <0.0001* |

**Table A3:** Statistical results of the follow-up Bonferroni tests with the number of cases associated with each avian migratory status. An asterisk (*) denotes that the p-value is statistically significant.

| Goodness of Fit Test Bonferroni Corrections: Distribution of Case Across Migratory Status, df = 1 |  |  |  |
| --- | --- | --- | --- |
| Variable One | Variable Two | $\chi^2$ value | *p-value* |
| Breeding | Non-breeding | 0.33333 | 0.5637 |
| Breeding | Year-round Residents | 177.09 | <0.0001* |
| Breeding | Unknown | 0.33333 | 0.5637 |
| Non-breeding | Year-round Residents | 180.02 | <0.0001* |
| Non-breeding | Unknown | 0 | 1 |
| Year-round Residents | Unknown | 180.02 | <0.0001* |

**Table A4:** Statistical results of the follow-up Bonferroni tests with the number of cases associated with each avian foraging preference. An asterisk (*) denotes that the p-value is statistically significant.

| Goodness of Fit Test Bonferroni Corrections: Distribution of Cases Across Foraging Preferences, df = 1 |  |  |  |
| --- | --- | --- | --- |
| Variable One | Variable Two | $\chi^2$ value | *p-value* |
| Birds | Insects | 17.789 | <0.0001* |
| Birds | Omnivores | 114.05 | <0.0001* |
| Birds | Seeds | 5.26 | 0.02181 |
| Birds | Small Animals | 35.714 | 0.05878 |
| Insects | Omnivores | 60.129 | <0.0001* |
| Insects | Seeds | 4.5918 | 0.03212 |
| Insects | Small Animals | 29.121 | <0.0001* |
| Omnivores | Seeds | 87.811 | <0.0001* |
| Omnivores | Small Animals | 128.03 | <0.0001* |
| Seeds | Small Animals | 14.222 | 0.0001* |

**Table A5:** Results of the follow-up Bonferroni tests with the number of cases associated with each avian habitat in our representative sample. An asterisk (*) denotes that the p-value is statistically significant.

| Goodness of Fit Test Bonferroni Corrections: Distribution of Cases Across Species Primary Habitat, df = 1 |  |  |  |
| --- | --- | --- | --- |
| Variable One | Variable Two | $\chi^2$ value | *p-value* |
| Forest | Open Woodlands | 19.93 | <0.0001* |
| Forest | Towns | 55.194 | <0.0001* |
| Open Woodlands | Towns | 10.922 | 0.00095* |

**Table A6:** Results of the bivariate Chi-Square Tests of Association for each variable permutation. An asterisk (*) denotes that the p-value is statistically significant.

| Bivariate Chi-Square Tests of Association for the Independent Variables, n = 187 cases |  |  |  |  |  |  |
| --- | --- | --- | --- | --- | --- | --- |
| Variable One | Variable Two | $\chi^2$ | *p-value* | $V_{\text{cramer}}$ | CI<sub>95%</sub> | Df |
| Migratory Status | Habitat | 3.87 | 0.69 | 0.00 | 0.00 – 1.00 | 6 |
| Migratory Status | Foraging Preference | 15.17 | 0.23 | 0.07 | 0.00 – 1.00 | 12 |
| Migratory Status | Foraging Niche | 139.83 | <0.0001* | 0.40 | 0.40 – 1.00 | 9 |
| Migratory Status | Natural Food: Birds | 7.70 | 0.05 | 0.16 | 0.00 – 1.00 | 3 |
| Migratory Status | Natural Food: Insects | 0.31 | 0.96 | 0.00 | 0.00 – 1.00 | 3 |
| Migratory Status | Natural Food: Seeds | 0.66 | 0.88 | 0.00 | 0.00 – 1.00 | 3 |
| Migratory Status | Natural Food: Berries and Plants | 3.92 | 0.27 | 0.07 | 0.00 – 1.00 | 3 |
| Habitat | Foraging Preference | 82.52 | <0.0001* | 0.45 | 0.33 – 1.00 | 8 |
| Habitat | Foraging Niche | 9.06 | 0.17 | 0.09 | 0.00 – 1.00 | 6 |
| Habitat | Natural Foods: Birds | 109.33 | <0.0001* | 0.76 | 0.63 – 1.00 | 2 |
| Habitat | Natural Foods: Insects | 13.9 | <0.0001* | 0.25 | 0.09 – 1.00 | 2 |
| Habitat | Natural Foods: Seeds | 20.47 | <0.0001* | 0.32 | 0.17 – 1.00 | 2 |
| Habitat | Natural Foods: Berries and Plants | 135.74 | <0.0001* | 0.85 | 0.72 – 1.00 | 2 |

**Table A6:** Results of the bivariate Chi-Square Tests of Association for each variable permutation. An asterisk (\*) denotes that the p-value is statistically significant. (*Continued*)
| Bivariate Chi-Square Tests of Association for the Independent Variables,<br>n = 187 cases |  |  |  |  |  |  |
| --- | --- | --- | --- | --- | --- | --- |
| Variable One | Variable Two | $\chi^2$ | *p-value* | $V_{\text{cramer}}$ | CI<sub>95%</sub> | Df |
| Foraging Preference | Foraging Niche | 201.66 | <0.0001* | 0.59 | 0.50 – 1.00 | 12 |
| Foraging Preference | Natural Food: Birds | 159.42 | <0.0001* | 0.91 | 0.78 – 1.00 | 4 |
| Foraging Preference | Natural Food: Insects | 108.64 | <0.0001* | 0.75 | 0.61 – 1.00 | 4 |
| Foraging Preference | Natural Food: Seeds | 103.98 | <0.0001* | 0.73 | 0.60 – 1.00 | 4 |
| Foraging Preference | Natural Food: Berries and Plants | 70.35 | <0.0001 | 0.60 | 0.46 – 1.00 | 4 |
| Natural Foods: Birds | Natural Foods: Insects | 6.95 | <0.0001* | 0.18 | 0.00 – 1.00 | 1 |
| Natural Foods: Birds | Natural Foods: Seeds | 27.73 | <0.0001* | 0.38 | 0.26 – 1.00 | 1 |
| Natural Foods: Birds | Natural Foods: Berries and Plants | 52.56 | <0.0001* | 0.53 | 0.40 – 1.00 | 1 |
| Natural Foods: Insects | Natural Foods: Seeds | 7.04 | <0.0001* | 0.18 | <0.0 – 1 | 1 |
| Natural Foods: Insects | Natural Foods: Berries and Plants | 7.62 | <0.0001* | 0.19 | 0.04 – 1.00 | 1 |
| Natural Foods: Seeds | Natural Foods: Berries and Plants | 18.88 | <0.0001* | 0.31 | 0.18 – 1.00 | 1 |
| Feederwatch | Habitat | 4.36 | 0.11 | 0.11 | 0.00 – 1.00 | 2 |
| Feederwatch | Foraging Preference | 133.29 | <0.0001* | 0.83 | 0.70 – 1.00 | 4 |
| Feederwatch | Foraging Niche | 157.48 | <0.0001* | 0.91 | 0.78 – 1.00 | 3 |
| Feederwatch | Natural Foods: Birds | <0.0001 | 0.97 | 0.00 | 0.00 – 1.00 | 1 |
| Feederwatch | Natural Foods: Insects | 30.27 | <0.0001* | 0.40 | 0.27 – 1.00 | 1 |
| Feederwatch | Natural Foods: Seeds | 27.77 | <0.0001* | 0.38 | 0.26 – 1.00 | 1 |
| Feederwatch | Natural Foods: Berries and Plants | 1.08 | 0.30 | 0.02 | 0.00 – 1.00 | 1 |
| Feederwatch | Migratory Status | 53.99 | <0.0001* | 0.52 | 0.39 – 1.00 | 3 |

